# Classifying Resting State Connectivity: Lag-One Autocorrelation and Pattern Differentiability

**DOI:** 10.1101/2025.06.17.659817

**Authors:** Yaohui Ding

## Abstract

Resting state functional connectivity (rsFC) and resting state effective connectivity (rsEC) are two of the most common measures that can be extracted from resting state functional magnetic resonance imaging (rs-fMRI) data. RSFC is often used to indicate the statistical dependencies among different brain regions of interest, whereas rsEC describes the causal influences among them. Many studies have explored utilities of rsFC and rsEC measures for classifying psychiatric conditions. Several studies showed that rsEC were better than rsFC features for classifying major depression (Frässle et al., 2020; Geng et al., 2018) and schizophrenia ((Brodersen et al., 2014)). However, no study to-date has investigated whether rsEC is *inherently* better than rsFC for classifying psychiatric conditions or the impact of autocorrelation on classifying rsFC, even though autocorrelation is known to be present in rs-fMRI data. To fill these gaps, we performed a series of computational experiments, by varying the size of the network and the number of participants, to gain some insight into these two aspects of supervised classification with resting state connectivity. Contrary to what has been reported in the literature, the results from our study suggest that rsEC cannot be, in principle, better than rsFC features for classification. In fact, rsEC measures led to systematically worse classification results, compared to rsFC features. In terms of the impact of autocorrelation, we found that lag-one autocorrelation could lead to both false negative and false positive classification results for studies with a small sample size.

## Introduction

The brain is an organ that exhibits both high levels of functional specialization and integration. Over the last few decades, there has been an increasing interest in studying functional integration of different brain regions at rest. Resting state functional magnetic imaging (rs-fMRI) is one of the most commonly used experimental paradigm for measuring brain activity at rest. Within this context, two broad categories of brain connectivity have been defined, namely resting state functional connectivity (rsFC) and resting state effective connectivity (rsEC). Resting state functional connectivity represents the statistical dependencies among different brain regions, whereas rsEC describes the causal influences among these brain regions (Friston, 2011). Many studies have explored the utilities of rsFC measures for classifying psychiatric conditions, e.g., major depression (Zeng et al., 2012), autism (Hull et al., 2017), and schizophrenia (Arbabshirani et al., 2013). In recent years, rsEC measures have also been utilized for classifying these psychiatric conditions. Furthermore, several studies showed that rsEC were better than rsFC features for classifying major depression ((Frässle et al., 2020; Geng et al., 2018) and schizophrenia ((Brodersen et al., 2014). However, no study to-date has investigated whether rsEC is inherently, i.e., from a theoretical perspective, better than rsFC for classifying psychiatric conditions. Additionally, no study to-date has examined the impact of autocorrelation on supervised classification with resting state functional connectivity, either, even though autocorrelation is known to be present in rs-fMRI data. The current computational modeling study is primarily concerned with two questions. First, does autocorrelation affect the classification results if we were to use resting state functional connectivity features for classifying psychiatric conditions? Second, is resting state effective connectivity inherently better than resting state functional connectivity for classifying psychiatric conditions?

### Resting State Functional Connectivity (rsFC)

There are many ways to define resting state functional connectivity (rsFC), seed-based, region of interest (ROI)-based, Independent Component Analysis (ICA), and graph theoretical approaches among others. In this study, we will adopt the ROI-based pairwise correlation approach of defining resting state functional connectivity. In the ROI-based approach, the fMRI time course from an ROI is correlated with fMRI time series from the rest of the brain. The main advantage of the ROI-based approach is that it is computationally simple thus can be easily extended to the whole brain (Lv et al., 2018). Additionally rsFC features from ROI-based functional connectivity analysis are easy to interpret, compared to results from the ICA approach. However, functional connectivity measures from both the seed-based and ICA approaches are correlational in nature, whereas effective connectivity measures from dynamic causal modeling are directional and causal in nature.

### Resting State Effective Connectivity (rsEC)

Resting state effective connectivity can be estimated from spectral dynamical causal modeling of the resting state fMRI data (Friston et al., 2014). Dynamic causal modeling (DCM) is a model-based and principled approach to inferring the hidden causal structure of distributed neural processes from observed neurophysiological data (Friston et al., 2003). Many variants of DCM have been developed over the years to extend the capabilities of the standard DCM approach. One of such extensions is the spectral dynamical causal modeling, which was developed for modeling resting-state fMRI data. Spectral DCM rests on a family of generative models of the complex cross spectra of the observed blood oxygenation level dependent (BOLD) signals and the inversion and comparison of these generative models via variational Bayes, Bayesian model reduction, and parametric empirical Bayes (Zeidman et al., 2019). However, the generative model in spectral DCM is more or less the same to the one in standard DCM (see Appendix-III for a detailed description of the generative model in DCM). Similarly, the model inversion and comparison procedures are also largely the same for these two approaches.

## Methods

### Experiment One: Starting from Two Distinct rsFC Patterns

For this computational experiment, we begin with the assumption that the rsFC patterns from n brain regions of interest for the clinical group (Figure-1.B-1) and healthy control group (Figure-1. B-2) are clearly distinct, which will serve as the templates for generating the rsFC matrices for the participants in each groups. Next, we generated *N*_1_ and *N*_2_ matrices for the clinical and healthy control groups, respectively, by sampling from two Wishart distributions. Then, we normalize these samples to get the correlational matrices (rsFC patterns) because samples from Wishart distributions are covariance matrices. Next, we use a lag-one vector autoregressive model to generate synthetic BOLD signals (Figure-1. C-2) from n ROIs for each participant, with subject- and ROI-specific lag-1 autocorrelation sampled from a random uniform (0.3, 0.8) distribution. The rsFC matrices are then re-calculated based on the synthetic BOLD signals, which has autocorrelation added. To get the rsEC features, we fit a spectral DCM model (Friston et al., 2014) to each participant’s BOLD data. Finally, we do supervised classification (support vector machine and logistic regression) with the rsFC features (before and after autocorrelation has been added) and the rsEC features to try to differentiate the clinical from healthy control group. The figure below illustrates the majors steps involved in this generative process using a 15-ROI network, see Appendix-I for a much more detailed description of all the steps involved.

**Figure-1.**
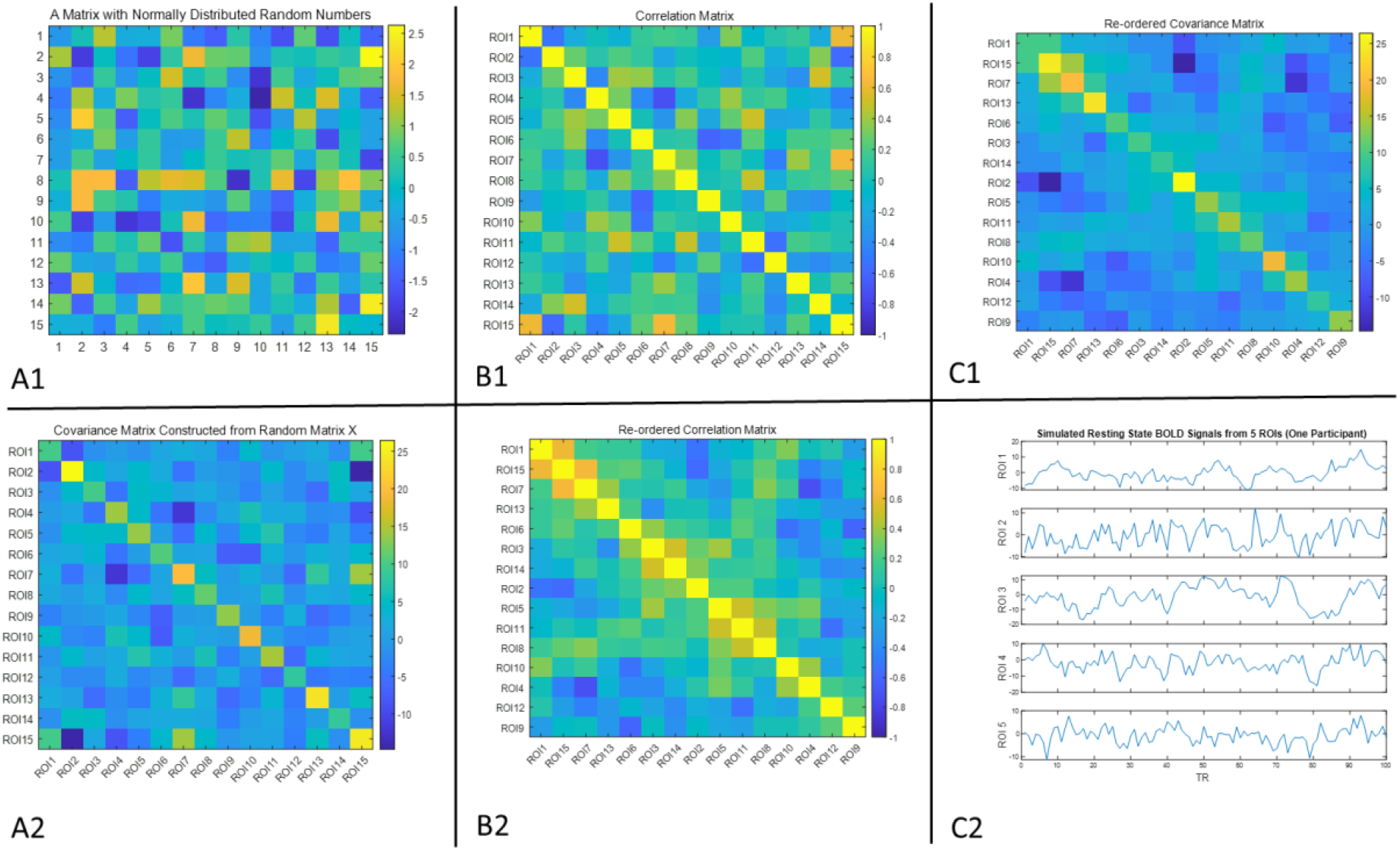
An illustration of the major steps involved in the data generation process.

### Experiment Two: Starting from two similar rsFC patterns

The procedure of experiment two is equivalent to that of experiment one, except that we start with the assumption that the rsFC patterns for the clinical and healthy control groups are very similar. Therefore, we should not expect to be able to distinguish the clinical from the healthy control group with the rsFC features.

## Results and Discussion

### The Impact of Lag-One Autocorrelation on Classifying Resting State Functional Connectivity

The results from our study suggest lag-one autocorrelation has a non-negligible effect on the classification results with rsFC features, when the size of the network and the size of the study are small (Figure-2. A1). For example, the classification accuracy from a medium-sized study (N=100) with rsFC features from a 10-ROI network could potentially be as low as 0.57, even though the classification accuracy is perfect with rsFC features without autocorrelation. Similarly, the classification accuracy from a small study (N=60) with rsFC features from a 50-ROI network could be as low as 0.72. However, the effect of lag-one autocorrelation does seem to level off as the size of the network increases (Figure-2. A1). This means that lag-one autocorrelation, if not corrected, could lead to false negatives for studies with small sample sizes and rsFC features extracted from small networks. In contrast, if we assume that the rsFC patterns from the clinical and healthy control groups are not distinguishable to begin with, we see that lag-one autocorrelation tends to have a small effect on the classification accuracy regardless the size of the network (Figure-2. A2). For example, the accuracy from a medium-size study with rsFC features extracted from a 50-ROI network is roughly 0.65, even though expected classification accuracy should be 0.5 (chance-level). This means that if autocorrelation is not corrected, it could also lead to false positive results in small and medium-sized studies. However, the effect of lag-one autocorrelation on classification accuracy does seem to be minimal for studies with a very large sample size (multi-site study; see the green line in Figure-2. A2).

**Figure-2.**
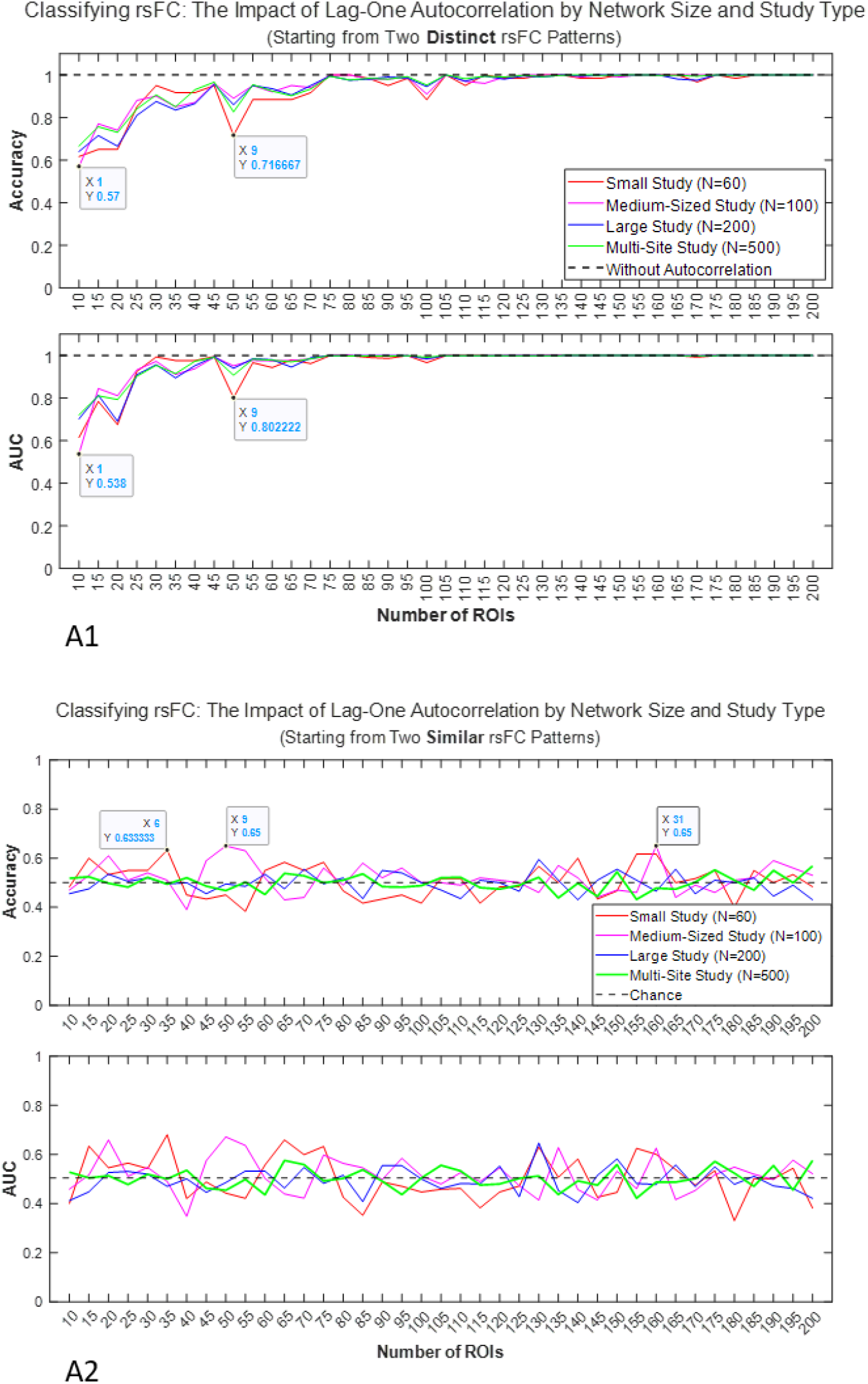
The Impact of Lag-One Autocorrelation on classifying rsFC Features

### RSEC is Not Principally Better than rsFC for Classifying Psychiatric Conditions

Contrary to what has been reported in the literature, the results from our study suggest that rsEC cannot be, in principle, better than rsFC features for classification. In fact, rsEC measures led to systematically worse classification results, compared to rsFC features. Starting from the assumption that the rsFC patterns for the clinical and healthy control groups are distinct, we see that rsEC features lead to much worse classification results (figure-3. A1). For example, the classification accuracies with rsEC features from 6-ROI, 12-ROI, and 20-ROI networks are 0.6, 0.74, and 0.61, respectively. This suggests that it may not be worth going through the trouble of estimating resting state effective connectivity, which is computationally expensive, if rsFC features led to good classification results. However, if one does not get good classification accuracy with rsFC features, it may not be worth the effort of estimating the rsEC features, either, because they would also lead to null results (see figure-3. A2).

**Figure-3.**
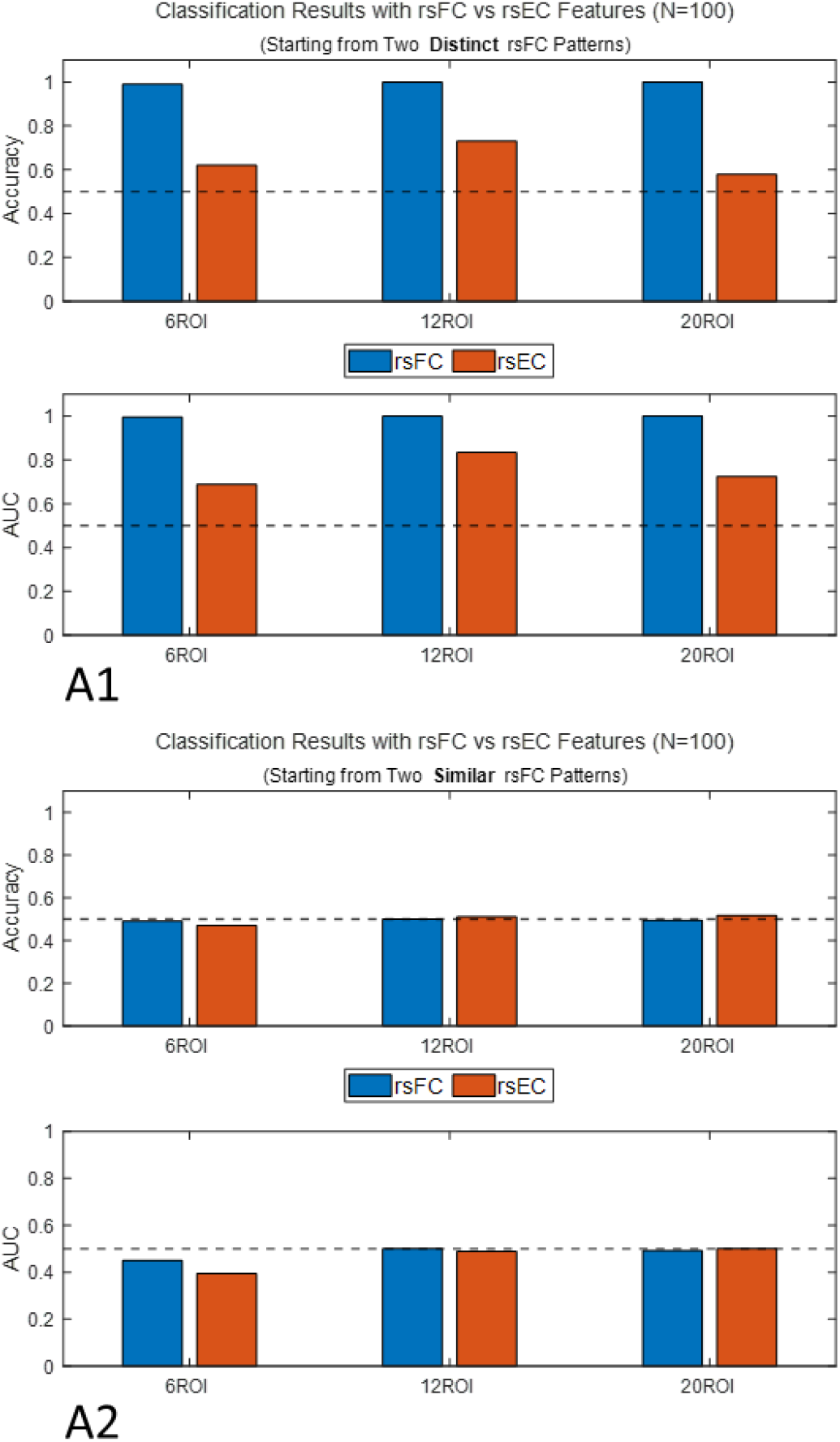
Comparing classification results with rsFC vs rsEC features

## Limitations and Future Directions

There are several limitations to the current computation modeling study. First, we assumed that resting state BOLD signals could be modeled with a lag-one autoregressive model. Though this is a reasonable assumption with some support in the literature (Liegeois et al., 2017), higher-order (lag-2 or lag-3) autocorrelation and cross-correlation may also need to be taken into account to model resting state fMRI data more accurately. The second major limitation of the current study is that we did not examine the impact of signal-to-noise ratio on classifying resting state functional and effective connectivity. Instead, we assumed that the noises in the BOLD signals have been sufficiently cleaned. This is a decision made for practical reason, because we were primarily concerned with examining the impact of autocorrelation on resting state functional connectivity and comparing the utilities of rsFC and rsEC features for classification. Additionally, we varied the size of the network and the number of participants for experiment one and two.

In terms of future directions, varying the amount of randomness that is added to the rsFC pattern at the subject level, which can be accomplished by changing the degrees of freedom parameter of the Wishart distribution in the data generation process, would also make the conclusions more robust. Finally, it would be hugely beneficial to test these ideas on empirical data, i.e., comparing classification results with and without autocorrelation corrected and with rsFC and rsEC features.

## Appendices

### Appendix-I: Data Generation Mechanism

**Step one**: Generate a 15 *by* 15 matrix *X* (full rank), where *x*_*ij*_*∼normal*(0,1), *i* = 1,2, …, 15 *and j* = 1,2, …,15.

**Step two**: Compute *A* = *X*^*T*^*X*. It can be shown that *A* is symmetric positive definite (SPD) if *X* is full rank. More importantly, it can also be shown that every SPD matrix is the covariance matrix of some random vector, up to some scaling constants. See Appendix-II for a detailed proof.

**Step three**: Compute the correlation matrix *Corr* = *sqrt*(*diag*(*A*))^−1^ * *A* * *sqrt*(*diag*(*A*))^−1^.

**Step four**: Do hierarchical clustering on the correlation matrix.

**Step five**: Re-order the covariance and correlation matrices. Do Cholesky decomposition on the re-ordered covariance and correlation matrices to check if they are still symmetric positive definite (they should be). Uses this re-ordered correlation as the template rsFC matrix for group one, e.g., the healthy control group.

**Step six**: Generate *N*1 matrices for **group one** using the template. Specifically, use the re-ordered covariance matrix as the scale matrix *V* of a Wishart distribution with 1000 degrees of freedom and generate *N*1 samples from this distribution *W*_*i*_*∼Wishart* (*V, nDOF*), where i = 1,2, … N1, and *nDOF* = 1000. Normalize these samples, 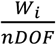. Additionally, compute the correlation matrices using the formula in step three.

**Step seven**: Generate *N*2 matrices for **group two** using the original (un-ordered) covariance matrix from step two as the template, following the same procedure described in step six.

**Step eight**: For each participant, generate synthetic BOLD signals using a lag-one vector autoregressive model, VAR(1).

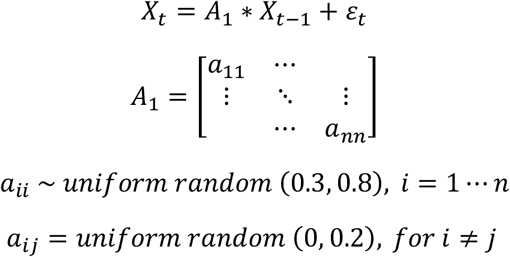

Where, *X*_*t*_ is a vector of BOLD signals from *n* ROI at time *t, A*_1_ the matrix of lag-1 autocorrelation and cross-correlation coefficients, and *ε*_*t*_ the residue signal which is sampled from a multivariate normal distribution, *N*(**0**, *W*_*i*_), using the sample covariance matrices generated in step six (**group one**) or step seven (**group two**).

**Step nine:** Re-compute the covariance and correlation matrices for each participant based on the synthetic BOLD signals generated in step eight, which has lag-one autocorrelation added.

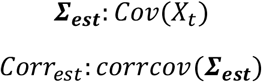

### Appendix-II: Every Symmetric Positive Definite Matrix is a Covariance Matrix

#### Proposition No. 1

A = X^T^X is symmetric positive definite if X is full rank.

Proof: Let X ∈ *R*^*n x n*^ be a matrix of samples from standard normal distribution.

1.1. Symmetry

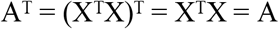

1.2. Positive Definiteness

Let v ∈ *R*^*n*^, and v ≠ 0. Consider the quadratic form:

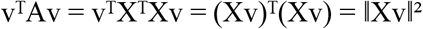

Since X has full column rank, Xv ≠ 0 for all v ≠ 0. Therefore:

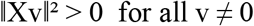

Hence:

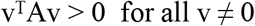

Therefore, A is symmetric positive definite (SPD).

#### Proposition No. 2

Every symmetric positive definite matrix A is the covariance matrix of some random vector z, up to some scaling constants.

Proof: Let A ∈ ℝ^pxp^ be a symmetric positive definite matrix.

2.1. Cholesky Decomposition of A

Since A is symmetric and positive definite, it has a Cholesky decomposition:

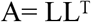

Where L is a lower triangular matrix with positive diagonal entries.

2.2. Construct a New Random Vector x

Suppose z ∈ ℝ^p^ is a random vector with zero mean and identity covariance matrix, i.e., z ∼ N (0, I). Define a new random vector x ∈ ℝ^p^ such that,

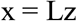

Then the mean of x is,

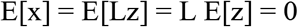

2.3. Compute the Covariance Matrix of x

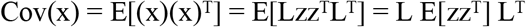

Since E[zz^T^] = I, we have:

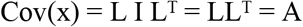

Therefore, matrix A is the covariance matrix of the random vector x = Lz.

### Appendix-III: The Generative Model in Dynamic Casual Modeling

#### The Neural State Equation

The original dynamical causal modeling (DCM) framework proposed in Friston et al. (2003) is a state space model that consists of a state transition equation that describes how the neural states evolve over time and a forward model that translates neural states into observed blood oxygenation level dependent signals. The neural state equation takes (1) on a bilinear form,

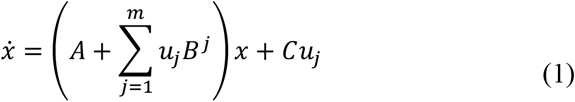

Where, *x* represents the neural state. Matrix *A* is the endogenous effective connectivity that describes the rates of change of neural activity of each brain region due to the neural activity from other brain regions in the system or the brain region itself (self-connection). Matrix *B* specifies how the endogenous effective connectivity matrix *A* is modulated by external inputs *u*, hence it is called modulatory effective connectivity. Finally, the rate of change in neural activity at each brain region due to the direct influence of the external inputs u is encoded by matrix *C*. Therefore, within the dynamic causal model framework, external inputs can have either (or both) modulatory effects that modulate the endogenous effective connectivity among different brain regions or driving effects that directly drive the neural responses at different brain regions. For resting state fMRI data where there are no external inputs (at least in the sense of experimentally controlled external inputs) to the brain, the neural state equation can be simplified into (2), where *w* is additive Gaussian noise.

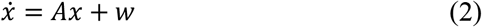

#### The Hemodynamic State Equations

The forward model consists of the hemodynamic state equations and the BOLD output nonlinearity. The hemodynamic state equations (3-6) relate the mean (and hidden) neural activity from each brain region to the hemodynamic states from that brain region. It is comprised of two parts. The first part of the hemodynamic observation equation describes how the neural activity *x* from a brain region induces a vasodilatory signal *s* and how *s* further induces a regional cerebral blood flow (*rCBF*) signal *f*. Together equations (3) and (4) describe the behavior a dampened oscilator with auto-regulatory feedback, where, *s* is the vasodilatory signal, *f* the *rCBF* flow signal, *κ* the rate constant of vasodilatory signal decay, and *γ* the rate constant for autoregulatory feedback by blood flow. The second part of the hemodynamic equation is the so-called Balloon model (Buxton et al., 1998), which describes how regional cerebral blood flow *f* leads to changes in blood volume *v* and deoxyhemoglobin content *q*, equations (5-6), where, *tao* is the transit time, *alpha* the Grubb’s vessel stiffness exponent, *E*_0_ the capillary resting net oxygen extraction fraction, and *E* the oxygen extraction function (Stephan et al., 2007).

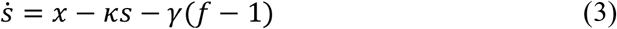

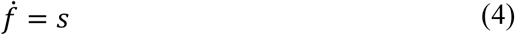

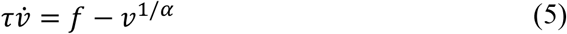

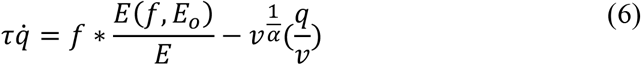

#### The BOLD Output Nonlinearity

Finally, the BOLD output nonlinearity describes how changes in *q* and *v* give rise to the BOLD signal that is detected with in an fMRI scanner.

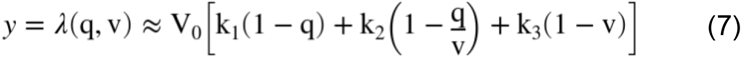

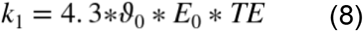

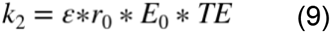

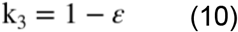

Where, *TE* is the echo time, *V*_0_ the resting venous volume, *E*_0_ the resting oxygen extraction fraction, *r*_0_ the slope of intravascular relaxation rate, *ε* the ratio of intra- to extra-vascular signal, *ϑ*_0_ the frequency offset (Stephan et al., 2007).

